# Sugar phosphate-mediated inhibition of peptidoglycan precursor synthesis

**DOI:** 10.1101/2024.11.13.623475

**Authors:** Megan R. Keller, Vijay Soni, Megan Brown, Kelly M. Rosch, Anas Saleh, Kyu Rhee, Tobias Doerr

## Abstract

Antibiotic tolerance, the widespread ability of diverse pathogenic bacteria to sustain viability in the presence of typically bactericidal antibiotics for extended time periods, is an understudied steppingstone towards antibiotic resistance. The Gram-negative pathogen *Vibrio cholerae*, the causative agent of cholera, is highly tolerant to β-lactam antibiotics. We previously found that the disruption of glycolysis, via deletion of *pgi* (*vc0374*, glucose-6-phosphate isomerase), resulted in significant cell wall damage and increased sensitivity towards β-lactam antibiotics. Here, we uncover the mechanism of this resulting damage. We find that glucose causes growth inhibition, partial lysis, and a damaged cell envelope in Δ*pgi*. Supplementation with N-acetylglucosamine, but not other carbon sources (either from upper glycolysis, TCA cycle intermediates, or cell wall precursors) restored growth, re-established antibiotic resistance towards β-lactams, and recovered cellular morphology of a *pgi* mutant exposed to glucose. Targeted metabolomics revealed the cell wall precursor synthetase enzyme GlmU (*vc2762*, coding for the bifunctional enzyme that converts glucosamine-1P to UDP-GlcNAc) as a critical bottleneck and mediator of glucose toxicity in Δ*pgi*. *In vitro* assays of GlmU revealed that sugar phosphates (primarily glucose-1-phosphate) inhibit the acetyltransferase activity of GlmU (likely competitively), resulting in compromised PG and LPS biosynthesis. These findings identify GlmU as a critical branchpoint enzyme between central metabolism and cell envelope integrity and reveal the molecular mechanism of Δ*pgi* glucose toxicity in *Vibrio cholerae*.

**Importance:** Sugar-phosphate toxicity is a well characterized phenomenon that is seen within diverse bacterial species, and yet the molecular underpinnings remain elusive. We previously discovered that disrupting *Vibrio cholerae’s* ability to eat glucose (by disrupting the *pgi* gene), also resulted in a damaged cell envelope. Upon deletion of *pgi*, glucose-phosphate levels rapidly build and inhibit the enzymatic activity of GlmU, a key step of bacterial peptidoglycan precursor synthesis. GlmU inhibition causes enhanced killing by antibiotics and a pronounced cell envelope defect. Thus, GlmU serves as a prime target for novel drug development. This research opens new routes through which central metabolism and sugar-phosphate toxicity modulate antibiotic susceptibility.

## Introduction

The antibiotic susceptibility spectrum, including resistance, tolerance, and persistence, continues to be a massive clinical threat (1, 2). Ranging from outright resistance, i.e. growth in the presence of antibiotics, to bacterial languishing (surviving and tolerating exposure antibiotics for a prolonged time), to having only a subset of the bacterial population persist and able to survive antibiotic exposure, there are many ways in which bacteria can respond to antibiotics (3–5). Furthermore, there has been an increase in studies finding that tolerance and persistence are steppingstones towards full resistance (6, 7). Discovering and understanding these midway points is needed to find novel potential antibiotic development routes, in order to solve this pressing issue facing our health care system.

With many common antibiotic development routes already being explored with varying degrees of success, interest in processes long overlooked (like tolerance) has recently surged. Particularly the contribution of central metabolic pathways to antibiotic susceptibility and infection outcomes have received considerable attention (8–13). Central carbon metabolism, for example, consists of interconnected pathways that ultimately produce energy for life as well as crucial precursors for biosynthesis of key macromolecules; most of these pathways are almost universally conserved (14–16). Glycolysis (more specifically, the Embden-Meyerhof-Parnas (EMP) pathway), the TCA cycle, and the pentose phosphate pathway are all vital carbon utilization networks found in virtually all extant species (17). Their interaction with many other cellular pathways (including cell envelope homeostasis) makes central carbon metabolism a potential central hub for determining antibiotic susceptibility, and consequently for developing novel forms of therapeutic intervention.

In a previous study, we discovered that deletion of *pgi*, a key enzyme in central carbon metabolism (EMP pathway) involved in the bidirectional conversion of glucose-6P to fructose-6P, causes cell wall damage and an increase in susceptibility to cell wall-acting antibiotics in the hypertolerant Gram-negative pathogen *Vibrio cholerae* (10). We found that these defects were associated with the intracellular accumulation of sugar phosphates and could be relieved by addition of the external cell wall precursor, N-acetylglucosamine (GlcNAc). Here, we sought to determine the molecular mechanism underlying sugar toxicity in the Δ*pgi* mutant. Genetic, metabolomic, and biochemical evidence suggest that glucose-1-phosphate inhibits GlmU function, thereby compromising the formation of UDP-GlcNAc, which results in inhibition of both peptidoglycan and potentially LPS biosynthesis. Our data thus identifies a new potential antibiotic target in *V. cholerae*, supporting the idea that metabolic disruptions could be weaponized to combat antibiotic tolerance and resistance.

## Results

### Glucose toxicity in a Δ*pgi* mutant manifests as morphological and functional damage to the cell envelope

To understand the negative effects caused by the addition of glucose in a Δ*pgi* mutant, we measured cellular morphology as well as survival in response to increasing glucose concentrations. WT, Δ*pgi*, and its complemented derivative were grown to exponential phase in M9 (minimal medium) supplemented with 0.2% casamino acids, followed by addition of increasing concentrations of glucose (0%, 0.02%, 0.2%, and 2%). Phase contrast microscopy after 3 hours of growth at 37°C revealed a notable morphology defect in Δ*pgi* (**Fig. 1A**), in essence recapitulating our previous observations in a more defined medium (10). An increase in glucose concentrations correlated with enhanced apparent cell death (visible as cell debris) and morphological defects, a typical response to inhibition of cell wall synthesis (18–21). To visualize cellular lysis further, we plated stationary phase cells on an LB agar plate with 0.2% glucose and 20 μg/mL of the cell impermeable β-galactosidase (LacZ) substrate CPRG (chlorophenol red-β-D-galactopyranoside) and incubated the plate overnight. Lysed cells will leak cytoplasmic LacZ into the medium and CPRG is hydrolyzed, resulting in a deep-red color change (22). While the Δ*pgi* mutant was able to form colonies on this plate, it demonstrated markedly enhanced cell lysis in the presence of glucose (**Fig. S1**). The combination of morphological defects and lysis suggests that both lipopolysaccharide (LPS) and peptidoglycan (PG) synthesis are at least partially affected in glucose-treated *pgi* mutants, as ordinarily PG synthesis inhibition results in spheroplast formation and little lysis in *V. cholerae*.

**Figure 1:**
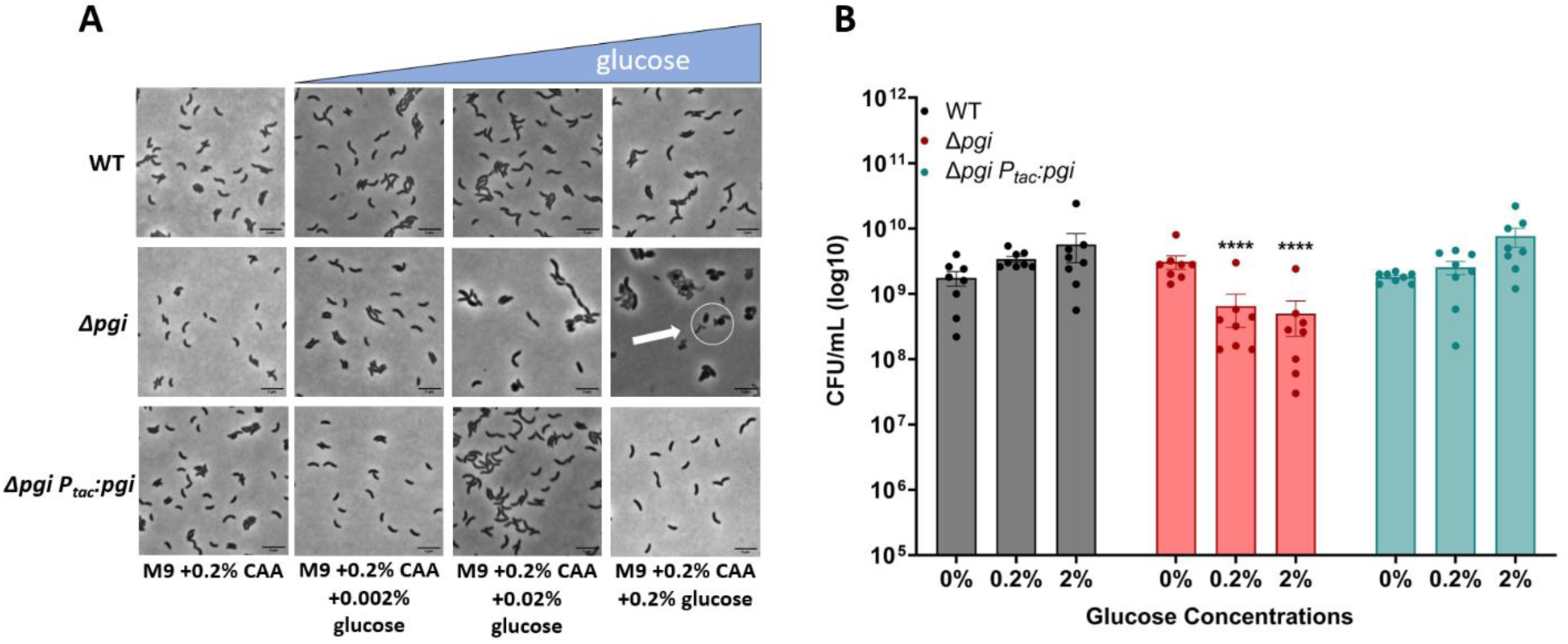
Glucose toxicity in Δ*pgi*. A) Cells were imaged after 3 hours of growth in the indicated conditions. Scale bar = 5 μm. White circle indicates aberrant cell morphologies B) CFU measurements from serially diluted cultures plated on M9 + 0.2% casamino acids agar after 3 hours of exposure to the indicated glucose concentrations. Mean with SEM plotted with 7 independent biological replicates. **=p<0.01, *** = p<0.001, ****=p<0.0001 (2-way ANOVA).

We next sought to assess the viability of Δ*pgi* cells in increasing glucose concentrations to determine the extent of glucose toxicity. Following the same experimental setup as described previously, increasing glucose concentrations were added to medium containing M9 + 0.2% casamino acids. Cells were incubated for 3 hours with defined glucose addition and then were serially diluted and plated on M9 agar containing 0.2% casamino acids. After an overnight incubation at 37°C, CFU/mL were counted. We noticed a slight but significant decrease in cell viability in a Δ*pgi* mutant that worsened with increasing glucose concentrations, reaching 10-fold at 2% glucose, while the WT remain unaffected (**Fig. 1B**). Collectively, these data indicate that glucose toxicity in a Δ*pgi* mutant results in a damaged cellular envelope.

### External N-Acetylglucosamine is sole carbon source to complement Δ*pgi* in glucose

We previously found that exogenous N-acetylglucosamine (GlcNAc) rescues Δ*pgi* defects in LB (10). However, since these experiments were done in LB (which produces a messy, poorly characterized physiology (23)), we sought to utilize a more chemically defined growth medium. In principle, the observed growth defect of Δ*pgi* in glucose could be due to either a decrease in cell wall precursor synthesis (which branches from glycolysis at fructose-6-phosphate, the product Pgi generates), or reduced flux into lower glycolysis, causing energy imbalance. Indeed, glucose phosphate toxicity (albeit in response to the glucose phosphate analog alpha-MG) in *E. coli* can be overcome by adding glycolytic intermediates, including fructose-6-phosphate (24). To test these ideas, we supplemented Δ*pgi* grown on M9 + 0.2% glucose with a panel of carbon sources, covering the spectrum of glycolysis and cell wall synthesis (**Fig. 2A**). After confirming cells could import and utilize these different carbon sources using a growth assay (**Fig. S2A**), we then plated serial dilutions of overnight cultures grown in M9 + 0.2% casamino acids, on M9 agar plates with glucose and the described carbon sources to assess rescue effects on the *pgi* mutant.

**Figure 2:**
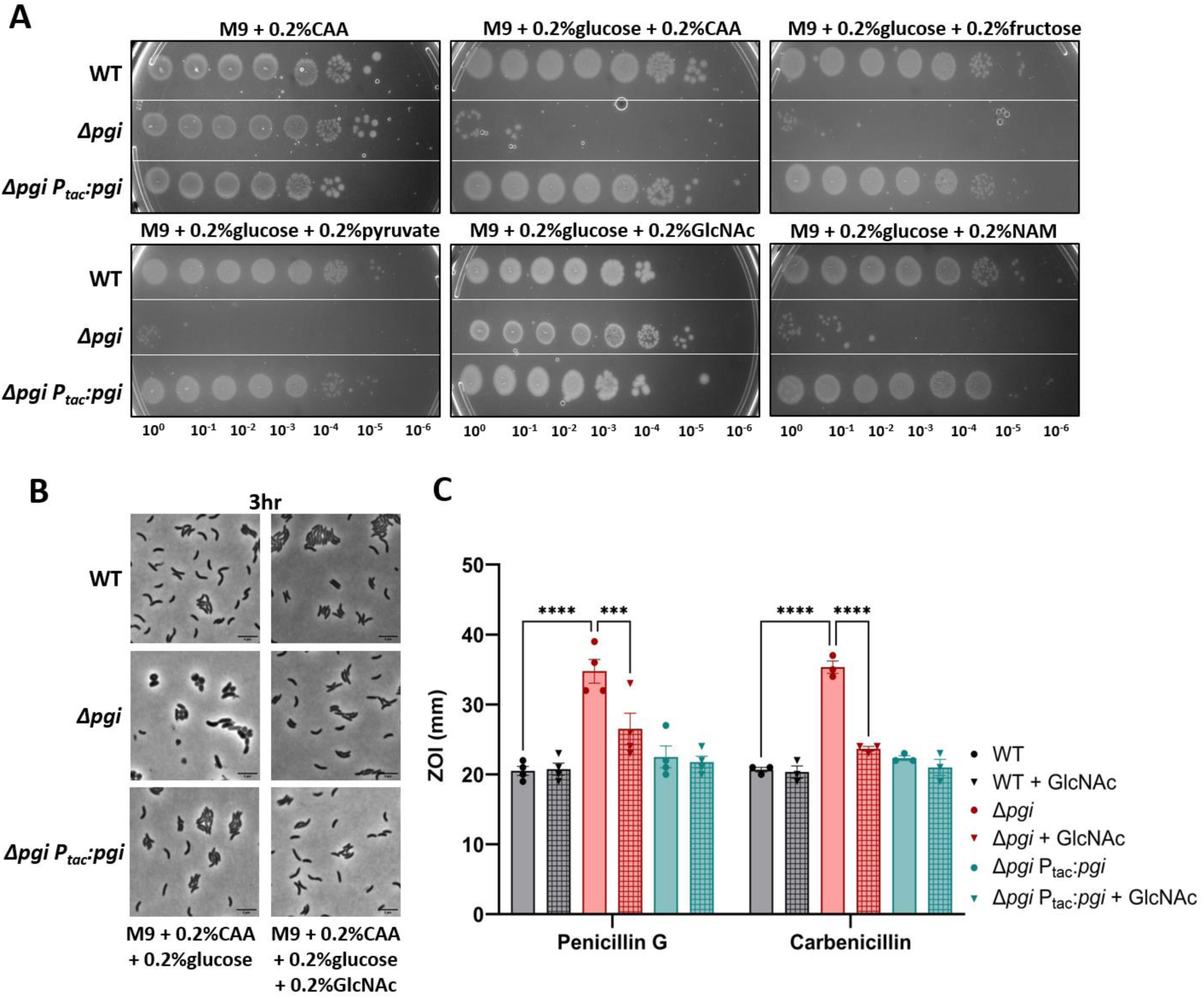
External GlcNAc is sole carbon source to complement Δ*pgi* in glucose. A) Serial dilutions of the indicated strains were plated on M9 minimal media supplemented with the indicated carbon sources and grown overnight at 37°C. B) The indicated mutant strains were imaged after 3 hours of growth at 37°C. Bar = 5 μm. C) Zone of inhibition measurements from a disk diffusion experiment on LB agar with or without the addition of 0.2% GlcNAc. Concentrations of the noted antibiotics are listed in Methods and Materials. Data represent at least 3 independent biological replicates; raw data points are shown with bars depicting mean with SEM. *** = p<0.001, ****=p<0.0001 (2-way ANOVA).

Interestingly, only GlcNAc could rescue Δ*pgi* growth and morphology in the presence of glucose (**Fig. 2A-B**). Neither fructose (which, upon import, gets converted to fructose-6-phosphate), pyruvate, nor succinate or glycerol rescued the Δ*pgi* growth defect on glucose, suggesting that Δ*pgi* glucose susceptibility is not primarily due to energy imbalance caused by disruption of glycolysis. We previously reported that a Δ*pgi* mutant was more sensitive to cell-wall targeting antibiotics (10). Therefore, we also sought to investigate the effect of GlcNAc on antibiotic susceptibility. Here, we added 0.2% GlcNAc to LB agar plates and measured the zone of inhibition in response to two β-lactam antibiotics (penicillin G and carbenicillin) (**Fig. 2C**). Addition of GlcNAc increased resistance to the Δ*pgi* mutant (though not to the WT). Thus, GlcNAc supplementation restores all characterized defects of a Δ*pgi* mutant.

Interestingly, MurNAc, another cell wall fragment, did not restore growth (**Fig. 2A**). This was curious, as both MurNAc and GlcNAc are cell wall precursors. MurNAc import occurs more upstream of *pgi* activity than GlcNAc and requires more steps to convert it into the common cell wall precursor glucosamine-6P. It is possible that the enzyme responsible for converting MurNAc-6P to GlcNAc-6P, MurQ, or the MurNAc transporter, MurP, are too inefficient for restoring optimal carbon flux, or not expressed under our growth conditions. Consistent with an inefficiency of MurNAc utilization, growth on MurNAc as sole carbon source resulted in much poorer yield than growth on GlcNAc (**Fig. S2**).

### Genetic manipulation of biosynthesis pathways reveals PG precursor demand in Δ*pgi*

We then began to genetically explore the GlcNAc import pathway and its connections between *pgi* and cell wall synthesis (summarized in **Fig. 3A**). We previously reported that glucose-6-phosphate accumulates in a *pgi* mutant; however, our untargeted metabolomics approach could not distinguish between glucose-6-phosphate (G6P) and glucose-1-phosphate (G1P). To dissect these two sugar phosphate species further, we constructed gene deletion and overexpression strains of the enzyme that converts G6P into G1P, *pgcA*, (*vc2095*), also known as *pgm* in *E. coli,* in a Δ*pgi* background. We hypothesized that reducing or boosting glucose-phosphate levels might mitigate or exacerbate *pgi* mutant phenotypes. We were additionally interested in the branch point enzymes NagB (*vca1025*) and GlmS (*vc0487*) for their role in regulating flux between glycolysis and cell-wall synthesis. We reasoned that by siphoning away early PG precursor metabolites (i.e. Glucosamine-6P) into glycolysis (via NagB), a *pgi* mutant would experience more defects, while GlmS overexpression should have the opposite effect. We thus tested these strains for their ability to modulate Δ*pgi* phenotypes. First, we tested PenG antibiotic susceptibility by measuring the zone of inhibition (**Fig. 3B**). *glmS* overexpression significantly reduced Δ*pgi* PenG sensitivity, suggesting that directing metabolic flux towards PG precursors and away from glycolysis was beneficial. Conversely, overexpression of *nagB* and *pgcA* tendentially enhanced PenG sensitivity, though this was not statistically significant (**Fig. 3A**).

**Figure 3:**
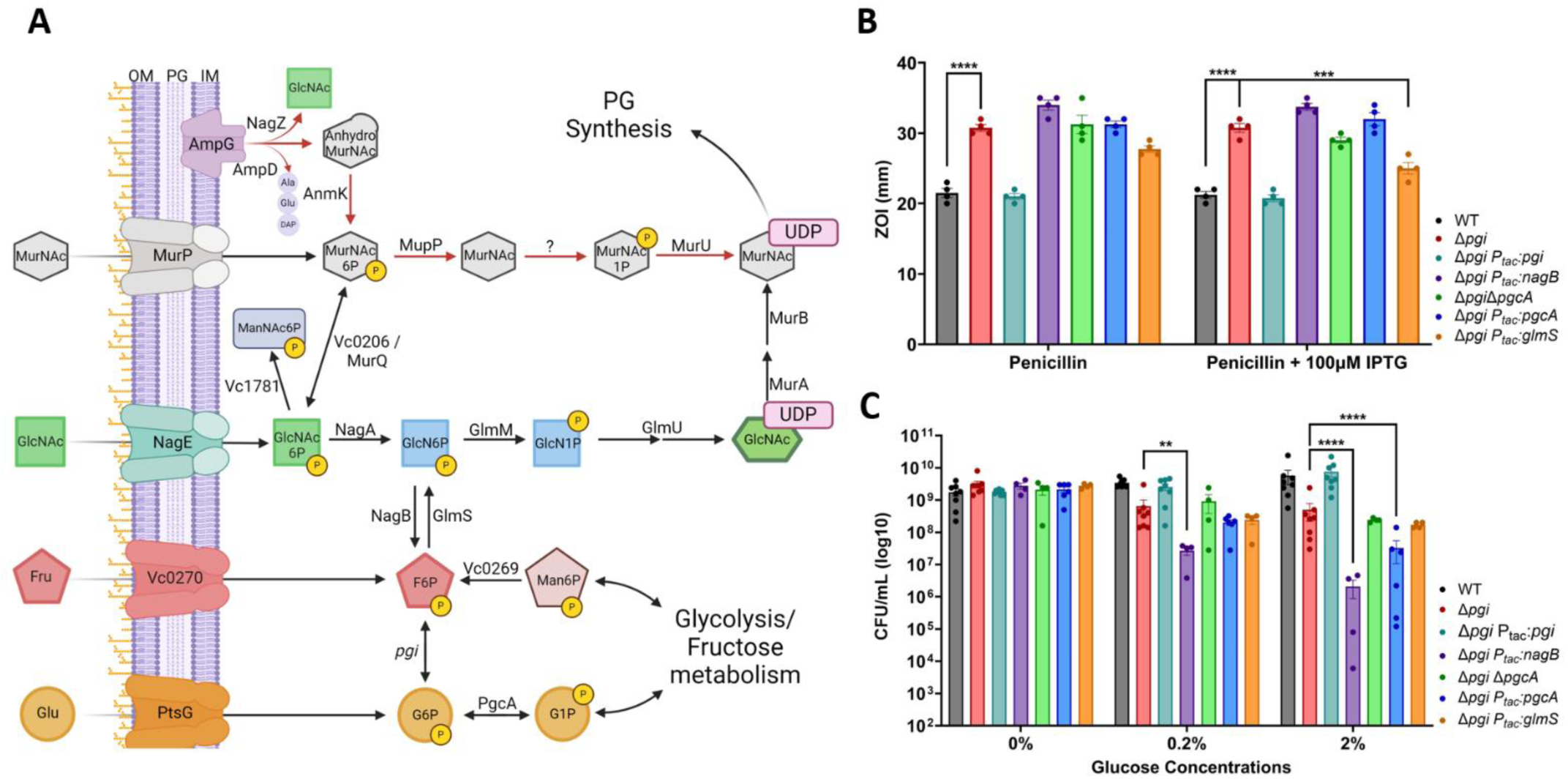
Genetic manipulation of biosynthesis pathways reveals an essentiality of glucosamine-6P in Δ*pgi*. A) A metabolic pathway diagram highlighting the PG recycling pathway, the conversion of glycolytic intermediates towards PG synthesis, and GlcNAc import. Glucose (Glu) can be imported through PtsG and readily converted to Glucose-6P. PgcA can also convert glucose-1P and glucose-6P interchangeably. Fructose (Fru) can be imported through *vc0270*, a member of the PTS system. Pgi interconverts G6P and F6P. Mannose-6P (Man6P) can be siphoned from central metabolism towards F6P. NagB and GlmS act as a bridge away from and towards PG synthesis, respectively. GlcNAc is imported through NagE and converted to Glucosamine-6P by NagA. GlmM and GlmU build the PG precursor UDP-GlcNAc, while MurA and MurB add some additional steps to create UDP-MurNAc. MurNAc can also be imported through MurP. As MurNAc-6P, this molecule can either be funneled back towards PG synthesis directly, through MupP, MurU and an uncharacterized enzyme, or recycled back into GlcNAc-6P by MurQ/*vc0206*. AmpG, a periplasmic PG fragment importer, can also supply internal GlcNAc from degraded cell wall products. B) Zone of inhibition data from treatment with PenG with and without overexpression induction. Statistical significance was evaluated via 2-way ANOVA from at least 5 independent replicates (raw data points shown). *** = p<0.001, ****=p<0.0001 C) Colony formation after 3 hours of glucose exposure with the indicated strains were plated on M9 + 0.2% CAA. Data for WT, Δ*pgi*, and complemented strain are reproduced from Fig.1 for comparison. Statistical significance was evaluated via log-transformed, 2-way ANOVA with 4 independent replicates (raw data points shown). *=p<0.1 **=p<0.01, *** = p<0.001, ****=p<0.0001.

Next, we turned to another phenotype, i.e. Δ*pgi*’s reduced survival in the presence of glucose. We thus treated the strains with either 0.2% or 2% glucose for 3 hours and then plated on M9 + 0.2% casamino acids (**Fig. 3C**). In both glucose concentrations, overexpressing *nagB* in a Δ*pgi* background resulted in significantly lower cell viability. At higher glucose levels, overexpression of *pgcA* in Δ*pgi* also became statistically significant in reducing cell viability. Collectively, these data point to a contribution of G1P in sugar phosphate toxicity of Δ*pgi and* suggest that carbon flux away from glycolysis helps this mutant, while flux away from cell wall precursor synthesis exacerbates its growth defect.

### Targeted metabolomics suggest a metabolite bottleneck around GlmU in glucose-treated *pgi* mutant cells

To further characterize the metabolic disruptions observed in a Δ*pgi* mutant and how the addition of GlcNAc could shift these metabolites, we conducted targeted metabolomics upon glucose exposure in M9 medium + CAA. Three hours after addition of either 0.2% glucose or a combination of 0.2% glucose and 0.2% GlcNAc, cells were pelleted, and metabolites were extracted using methanol. Samples were analyzed using LC-MS (see Methods and Materials), and peaks were compared to pure chemical standards. Upon normalizing the data to the casamino acid conditions, we noted a sharp increase in glucose-1P and glucose-6P, consistent with the *pgi* mutant’s inability to metabolize glucose through the EMP glycolysis pathway. The WT, but not Δ*pgi*, experienced an increase in pyruvate levels upon glucose addition. The combination of increased pyruvate and reduced G6P/G1P levels (and indeed very low F6P levels) in the WT perhaps indicates highly efficient upper glycolysis, which encounters a bottleneck at pyruvate processing. The relative lack of increase in pyruvate upon glucose addition in Δ*pgi* suggests that non-EMP pathways for glucose utilization (e.g., pentose phosphate pathway), and/or gluconeogenesis, may be inefficient in *V. cholerae,* at least under the conditions tested here. We also observed a sharp rise in glucosamine-1P levels (16-fold change), and a sharp, 36-fold decrease in UDP-GlcNAc, when Δ*pgi* was grown in medium containing glucose, compared to WT (**Fig. 4A**). Thus, the described sugar toxicity likely results from the inhibition of the enzyme converting glucosamine-1P to UDP-GlcNAc, which is GlmU (**Fig. 4B**). This bottleneck was slightly improved upon the addition of GlcNAc to the growth medium, suggesting GlcNAc-mediated relief of this inhibition. As expected from the pathway metabolizing external GlcNAc (**Fig. 3B**), the addition correlated with a slight increase in glucosamine-6P (into which external GlcNAc is converted by *V. cholerae* through the action of the NagE transporter and NagA deacetylase) (25).

**Figure 4:**
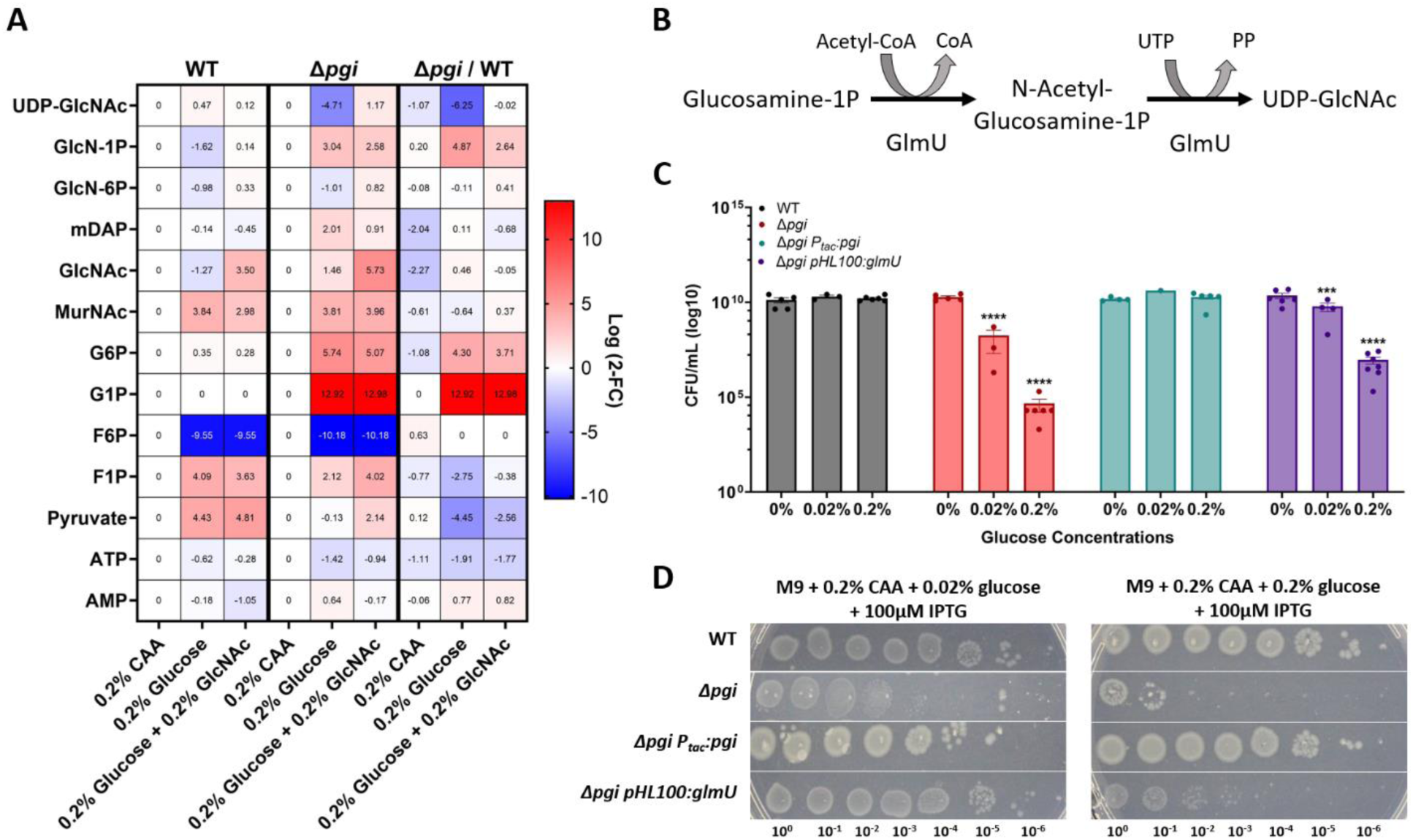
Targeted metabolomics of Δ*pgi* reveal bottleneck around GlmU activity. A) Heatmaps normalized to strain specific casamino acids conditions. The right column represents the change between strains in the indicated conditions. Log2 fold change is shown on the right side of the maps. B) The enzymatic reaction of GlmU. The acetyltransferase activity catalyzes N-acetylglucosamine-1P from glucosamine-1P and Acetyl-CoA. The second step is the uridyltransferase reaction which adds UDP onto GlcNAc, forming the end-product UDP-GlcNAc. C) Overnight cultures were serially diluted and spot-plated on MM agar with the indicated additions of percent glucose and 200uM IPTG. At least 4 independent replicates are presented, with raw data points and SEM. *** = p<0.001, ****=p<0.0001 (2-way ANOVA). Δ*pgi* was compared against WT values and Δ*pgi* pHL100: *glmU* was compared against Δ*pgi.* D) The indicated strains were grown overnight in M9 + casamino acids and the diluted and spot-plated on M9 agar plates containing CAA and the indicated glucose concentration.

Plausibly, the accumulation of glucose-1P and/or glucose-6P in the Δ*pgi* mutant competitively inhibits GlmU, downregulating cell wall synthesis. There is evidence of a similar effect in *Mycobacterium tuberculosis*, where at least glucose-1P competitively inhibits GlmU *in vitro* (26). Glucose-1P levels did not change (**Fig. 4A**) upon addition of GlcNAc, which may indicate that GlcNAc addition does not stop sugar phosphate accumulation, but rather circumvents PG precursor synthesis inhibition by supplying more substrate (which would suggest competitive inhibition). Fructose-6P levels in both WT and Δ*pgi* cells were significantly lower in glucose-containing medium. It is possible that flux into the TCA cycle and downstream respiration is faster in the presence of glucose, which causes a net relative decrease in F6P levels relative to growth in casamino acids (gluconeogenic conditions).

If GlmU is competitively inhibited by glucose phosphates, it should be possible to circumvent this inhibition by increasing the abundance of GlmU. Therefore, we next sought to genetically explore the role of GlmU by creating an overexpression construct in a Δ*pgi* background. Overexpression of *glmU* from a high copy number plasmid (see Methods and Materials), caused a significant increase in cell viability in the presence of glucose in comparison to Δ*pgi* alone (**Fig. 4C**). While there still was a decrease in overall survival, even when expressing *glmU* excessively, this can be explained by the glucose phosphate inhibition logic. We previously measured an over 200x increase in G1P/G6P in Δ*pgi* when grown in LB (10); adding a few more GlmU molecules is likely not sufficient to compensate for the intracellular flooding of G1P/G6P, especially when grown in medium with high glucose concentrations. We also overexpressed *glmU* from an overnight culture and plated serial dilutions on M9 + 0.2% CAA supplemented with either 0.02% glucose or 0.2% glucose (**Fig. 4D**). GlmU overexpression significantly improved Δ*pgi* plating efficiency on glucose, particularly at the lower glucose concentration, again supporting the idea of GlmU being the target of glucose toxicity in the Δ*pgi* mutant.

### *In vitro* biochemical validation of GlmU inhibition by glucose-1P

We next conducted an *in vitro* biochemistry assay using purified *V. cholerae* GlmU, to test whether G1P/G6P inhibition was direct. To this end, we tested a panel of G1P and G6P concentrations designed to mimic the glucose concentrations previously measured in Δ*pgi* (10) in a biochemical assay containing purified GlmU, as well as the reactants Acetyl-CoA, glucosamine-1-phosphate and UTP. After 30 min at 30°C, the reaction was stopped and analyzed for the emergence of the product of the GlmU reaction, UDP-GlcNAc, using LC-MS. Upon addition of increasing concentrations of G1P, but not G6P (**Fig. S3A**) to the reaction mix, we observed a significant decrease in UDP-GlcNAc abundance starting at 31.25mM of G1P, and near complete inhibition at 250mM (**Fig. 5C**). We also measured the intermediate product of GlmU’s bifunctional reaction, GlcNAc-1P. We found that at higher concentrations of G1P, GlcNAc-1P was also significantly reduced, suggesting that it is the acetyl-transferase activity of GlmU that is primarily affected by G1P inhibition.

**Figure 5:**
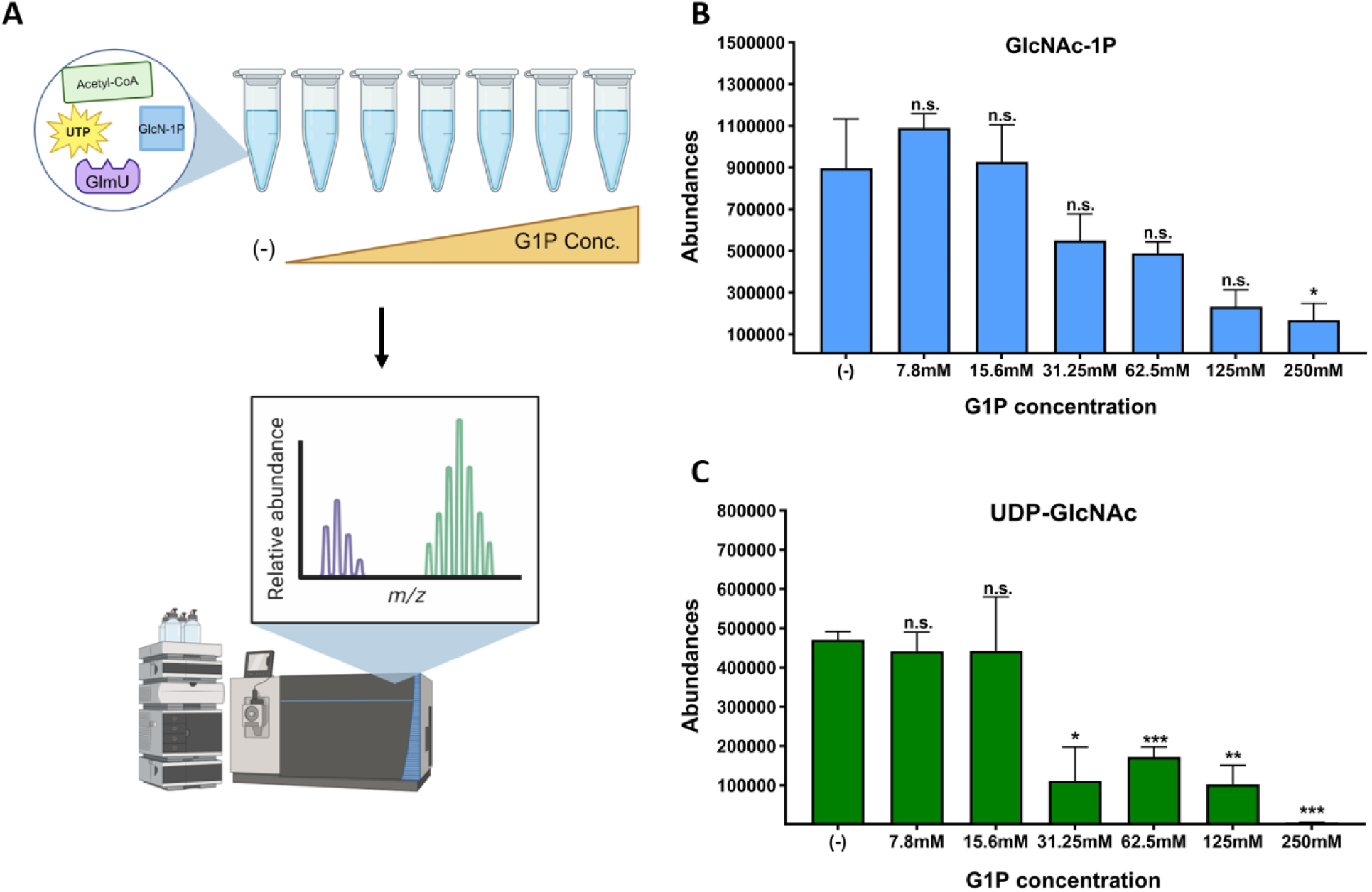
Glucose-1P inhibits GlmU biochemical reaction in a concentration dependent manner. A) Schematic portraying the *in vitro* biochemistry experimental design. Biochemical reaction components, UTP, Acetyl-CoA, GlcN-1P and GlmU, mixed with buffer, were added to test tubes. After incubation, the reaction was stopped and ran on the LC-MS machine. Abundance peaks were measured for the stated molecules. B) Peak values for GlcNAc-1P product from reactions with increasing glucose-1P concentrations. SEM plotted with 3 replicates. * = p < 0.05 (unpaired t-tests). C) Peak values for UDP-GlcNAc product from reactions with increasing glucose-1P concentrations (in mM, X-axis). Averages and SEM plotted from 3 replicates. * = p < 0.02, ** = p < 0.002, *** = p < 0.0005 (unpaired t-test).

### Molecular modeling reveals putative target site for glucose phosphate inhibition

We next sought to model the observed inhibition of glucose-1P against GlmU. While Alphafold3 is a powerful and accessible tool for modeling protein-ligand interactions, it is not well suited for our study as it is unable to model glucose-1P (27). Instead, we turned to Chai Discovery, a new online modeling program that predicts both protein multimers and protein-ligand binding (28). We obtained multiple models of predicted binding sites for both GlmU’s natural substrate, glucosamine-1P, but also its presumed inhibitor, glucose-1P, within the trimeric form of GlmU. Visualization of these molecular models revealed that glucosamine-1-P engages in polar interactions with Arg330, Lys348, Tyr363, Asn374, Asn383, and Lys389 (**Fig. 6A**). This model returned pTM and ipTM scores of 0.9556 and 0.9388, respectively; pTM scores > 0.5 and ipTM scores > 0.8 are considered confident predictions (29). Sequence alignments with GlmU^EC^ and GlmU^Mtb^, which have well characterized active sites (26, 30), indicate that the modeled residues likely form the acetyltransferase active site in GlmU^VC^ (**Fig. S5A-C**). Additionally, structural alignments of GlmU^VC^ to GlmU^EC^ and GlmU^Mtb^ indicate that the structure of GlmU is highly conserved across diverse organisms. Molecular modeling of the putative interaction between GlmU and glucose-1-P revealed a binding site at the same location as the natural substrate, indicating that glucose-1-P may competitively inhibit GlmU (**Fig. 6A**). This model returned similarly high pTM and ipTM scores (0.9549 and 0.9376 respectively). While there were some predictions that modeled G1P within the uridyltransferase pocket, the strength of the polar interactions were not as robust as the models that predicted G1P to bind within the acetyltransferase pocket (**Fig. S4B**). We additionally examined the predicted binding with glucose-6P, and found while it does bind in a similar fashion as glucose-1P, the pTM and ipTM scores were lower (**Fig.S4C-D**). How the phosphate location on the glucose molecule elicits such a drastically different inhibition response and binding affinity, remains to be explored. Together, this strongly suggests that G1P accumulation competitively inhibits the acetyltransferase activity of GlmU.

**Figure 6:**
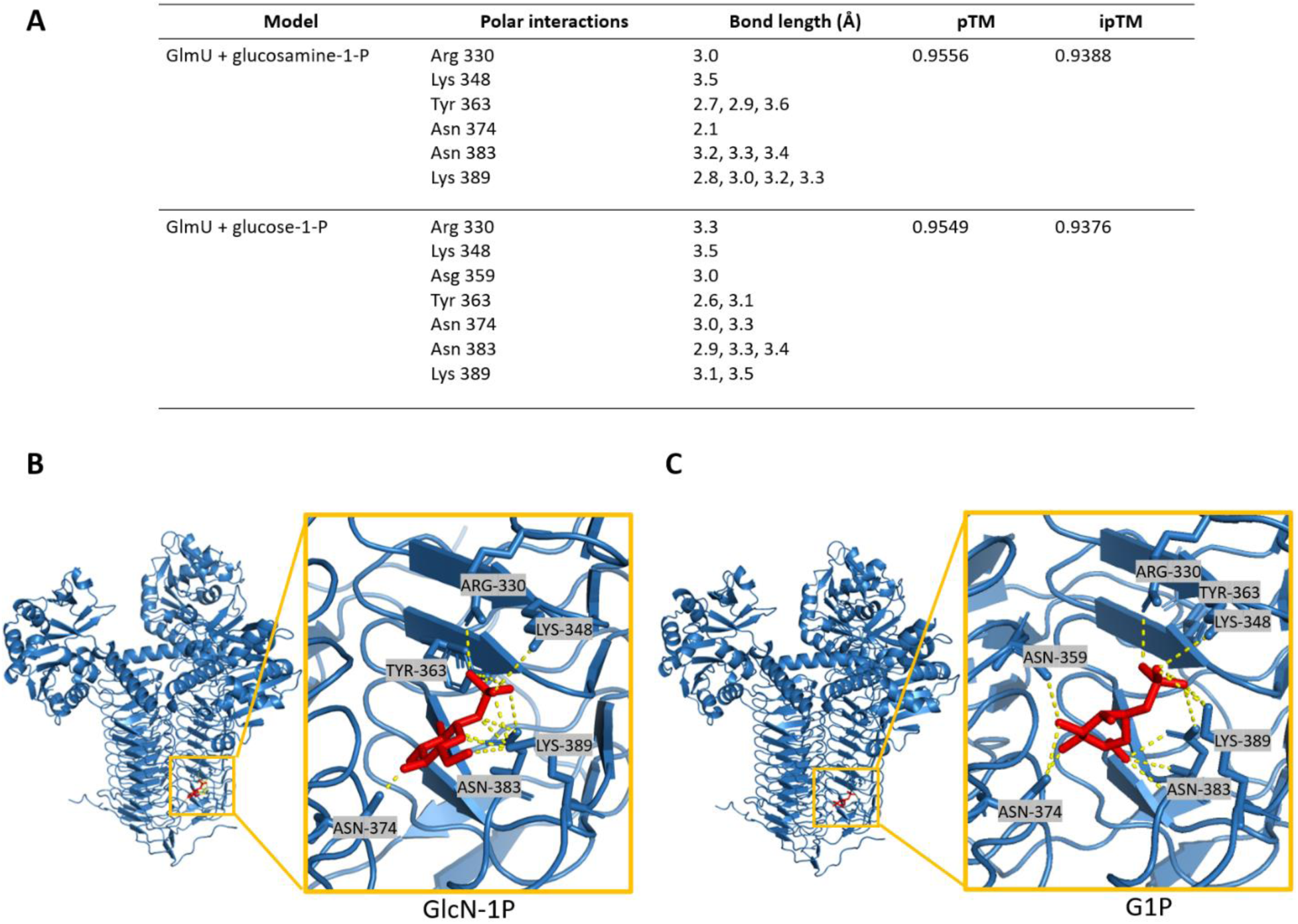
Molecular modeling reveals putative target site for glucose phosphate inhibition. A) Residues predicted to interact with glucosamine-1-P and glucose-1-P, Predicted Template Modeling (pTM) and Interface Predicted Template Modeling (ipTM). B) Molecular modeling of GlmU binding glucosamine-1-P (red) and the associated polar interactions. C) Molecular modeling of GlmU binding glucose-1-P (red) and the associated polar interactions.

## Discussion

Sugar-phosphate stress and its associated cellular defects remain underexplored in the context of antibiotic susceptibility. Here, we present data that elucidate the mechanism of glucose poisoning in *Vibrio cholerae*. We previously found that *pgi* mutation results in pronounced cell wall damage and concomitant increase in susceptibility to β-lactam antibiotics (10). In this study, we show that glucose toxicity in Δ*pgi* is due to (likely competitive) inhibition of GlmU (a key step in PG precursor synthesis) by sugar phosphate species. Sugar-phosphate toxicity has been studied for the past 7 decades (31–36), yet the mechanisms for glucose-related toxicity appear to be diverse, species-dependent, and poorly-understood (37, 38). In *B. subtilis,* a mutant defective in both glycolysis and pentose phosphate pathway builds up excessive G1P, which was suggested to inhibit an early PG precursor step, resulting in cell lysis (34). However, this observation was never followed up mechanistically. *E. coli* strains with a defective *pgi* experience significant sugar-phosphate stress, resulting in post-transcriptional regulation of *ptsG* to reduce sugar intake (38). While G6P levels are elevated in *Δpgi* backgrounds in *E. coli*, there is no observable cell-wall damage, and toxicity appears to be due to diversion of resources from glycolysis, rewiring of metabolism and possibly redox imbalance stress (35, 36, 39–41). The reduced glucose phosphate toxicity *in E. coli* may be due to a more enhanced flux into the pentose-phosphate pathway in this species, which could in principle efficiently remove G6P (41). Additionally, *E. coli* is known to have a robust glucose-phosphate stress response system regulated by small RNA molecule SgrS and its activator protein, SgrR (24, 42, 43). SgrS controls the excessive import of glucose through the PTS system by modulating *ptsG*. While *V. cholerae* encodes an SgrR homolog, the small RNA SgrS has not been identified.

Our data clearly show that sugar phosphate toxicity can directly contribute to cell envelope defects. While we do not have direct evidence of the specific mode of inhibition, our genetic, biochemical and modeling data point to competitive inhibition of GlmU by G1P, which could be explained by the similarity in structure between G1P and GlmU’s natural substrate GlcN-1P (**Fig. 7**). Additionally, if G1P is targeting the acetyltransferase domain, the alleviation experienced by addition of GlcNAc makes sense, as the external GlcNAc would readily be converted to first GlcN6-P and then GlcN-1P. More GlcN-1P would outcompete the G1P and restore GlmU functionality.

These findings more broadly shed light on the importance of central metabolism as a potential source of novel antibiotic targets. GlmU is a well conserved protein among highly relevant pathogens, most notably *Mycobacterium tuberculosis*. There have been extensive studies exploring the potential for GlmU as an anti-TB drug target, but little has been explored in other bacteria. Some studies have identified high throughput methods and computational models for drug screening against GlmU (44–49), while others have investigated the effect of depleting GlmU in infection models, mimicking the potential effects an inhibitor might have (50–55). In principle, UDP-GlcNAc biosynthesis serves as an ideal drug target, as it is required for not only PG, but also LPS biosynthesis. By designing targets for novel antibiotics that disrupt more than one biochemical pathway, resistance and mutations leading to reduced efficiency are less likely to occur.

## Methods and Materials

### Bacterial Strains and Growth Conditions

All *V. cholerae* strains used in this study are derivatives of *V. cholerae* El Tor strain N16961 and summarized in Table S1. *V. cholerae* was grown on Luria-Bertani (lysogeny broth) (LB) medium (for a 1 L bottle, 10 g Casein peptone, 5 g yeast extract, 10 g NaCl, and 12 g agar, all from Fischer Bioreagents) at 30°C or in M9 minimal medium (for a 1 L bottle, 15 g agar, 200 mL 5x M9 salts (for a 1 L bottle, 35 g Na 2 HPO 4 •7H 2 O, 15 g KH 2 PO 4, 2.5 g NaCl, 5 g NH 4 Cl), 0.5 mL 1M MgSO 4, 0.1 mL 1M CaCl 2, and 1 mL FeCl 3 /citric acid) at 37°C; 200 μg/mL of streptomycin was also added (N16961 is streptomycin resistant). Where applicable, growth media were supplemented with 0.2% glucose (w/v), 0.2% casamino acids (w/v), or 0.2% GlcNAc (w/v). All other carbon sources were also 0.2% (w/v).

For growth dynamic experiments, overnight cultures were diluted 100-fold into 1 mL growth media + streptomycin. 200μL of this seed stock were added to wells in a 100-well honeycomb and incubated in a Bioscreen growth plate reader (Growth Curves America) at 37°C with random shaking at maximum amplitude, and OD 600 recorded at 10 min intervals.

### Plasmid and strain construction

Oligonucleotides used in this study are summarized in Table S2. *E. coli* MFDλpir (a diaminopimelic acid [DAP] auxotroph) or SM10 λpir was used for conjugation into *V. cholerae*, for gene deletions and overexpression plasmids respectively (56). Overexpression strains were created using the chromosomal integration plasmid pTD101, a derivative of pJL1 containing lacIq and a multiple-cloning site under the control of the IPTG (isopropyl-β-d-thiogalactopyranoside)-inducible Ptac promoter, or pHL100mob, a non-integrative high copy number plasmid also inducible through IPTG(57). pTD101 integrates into the native *V. cholerae* lacZ (*vc2338*) locus. Genes for complementation experiments were amplified from N16961 genomic DNA (58), introducing a strong consensus ribosome-binding site (RBS) (AGGAGA), and cloned using Gibson assembly. Plasmids were colony PCR verified using primers 1 and 2 (pTD101) or 5 and 6 (pHL100mob). Gene deletions were constructed using the pTOX5 cmR/msqR allelic exchange system (59). In short, 500-bp regions flanking the gene to be deleted were amplified from N16961 genomic DNA by PCR and cloned into the suicide vector using Gibson assembly (60). Plasmids were colony PCR verified using primers 3 and 4. All plasmids were Sanger sequence verified before conjugation.

Conjugation into *V. cholerae* was performed by mixing overnight cultures 1:1 (100 μL donor plus 100 μL recipient) in 800 μL fresh LB, followed by pelleting (7,000 rpm, 2 min) and resuspending in 100 μL LB. The mixture was then spotted onto LB agar (with 600 μM DAP for E. coli MFDλpir growth) and incubated for 4 hr (overnight for pTOX5 deletions) at 37°C. Selection for single-crossover mutants was then achieved by streaking the mating mixture on either streptomycin (200 μg/mL) plus carbenicillin (100 μg/mL) (pTD101), streptomycin (200 μg/mL) plus kanamycin (50 μg/mL) (pHL100mob), or streptomycin (200 μg/mL) plus chloramphenicol (100 μg/mL) and 600 μM DAP for pTOX5 and incubating overnight at 37°C.

For pTD101 insertion, carbenicillin-resistant mutants were counterselected on salt-free LB supplemented with 10% sucrose and X-Gal (5-bromo-4-chloro-3-indolyl-β-d-galactopyranoside) (120μg/mL) and grown at ambient temperature for two days. White colonies (indicating a disrupted lacZ) were isolated, and PCR-verified using primers 23 and 24. For pTox-mediated recombination, chloramphenicol-resistant colonies were counter selected on M9 minimal medium containing 2% (vol/vol) rhamnose at 30°C for 18 hr. Deletions were verified by PCR using flanking and internal primers and verified with whole-genome sequencing.

### Cell Viability Assay and Glucose Time-dependent Killing Assay

To test cell viability, overnight cultures were added to sterile 1X PBS for serial dilution from 1:10 to 1:10^7^. 5 μL of overnight cultures and diluted cultures were spotted for CFU/mL on different media plates, as described in the figure legend. Dried plates were then incubated at 37°C (M9 agar) overnight and counted the next day. For glucose concentration-dependent experiments, strains were grown overnight in M9 + 0.2% casamino acids at 37°C. The following day, the cultures were diluted 1:1000 into fresh M9 + 0.2% casamino acids and incubated at 37°C for 3 hours. Then various concentrations of glucose were added to the media and incubated for another 3 hours at 37°C, then serially diluted onto M9 agar + 0.2% casamino acids and left overnight at 37°C. CFU/mL were counted the next day. For microscopy, strains were grown as previously described then imaged without fixation on M9 + 0.8% agarose pads using a Leica DMi8 inverted microscope.

### Antibiotic Sensitivity Assay

For zone of inhibition assays, a lawn of overnight cultures (100 μL) was spread on an LB agar plate with or without 0.2% GlcNAc and allowed to dry for 15 min. 10 μL of antibiotic solutions (100 mg/mL PenG or 100mg/mL carbenicillin) were placed on Thermo Scientific Oxoid Antimicrobial Susceptibility Test filter disks (6 mm, product code: 10609174) onto the agar surface and incubated at 30°C overnight before measurements.

### Metabolomics

Three biological replicates were grown in M9 + 0.2% casamino acids overnight at 37°C. 1mL of culture was pelleted (2 min, 7000 rpm) and washed with M9 media. The cultures were then added 1:50 into 5mL of M9 + 0.2% casamino acids and incubated at 37°C for 3 hours. Following incubation, either 0.2% glucose or 0.2% glucose and 0.2% GlcNAc were added to the tubes and incubated for an additional hour at 37°C. After treatment, 2 mL of sample were taken per condition and pelleted at 7000 rpm for 2 min. The supernatant was removed, and the cell material pellet was flash frozen with liquid nitrogen. 200 μL of cold 80% methanol was then added to the pellets. Pellets were stored at −80°C. These pellets were lysed and 3 µl samples were analyzed using Agilent InfinityLab Poroshell 120 HILIC-Z (Agilent 683775-924). The chromatographic separation employed two solvent phases: Solvent A (water + 10 mM NH4OAc + 5 mM InfinityLab Deactivator Additive, pH 9, adjusted with NH4OH) and Solvent B (85% ACN + 10 mM NH4OAc + 5 mM InfinityLab Deactivator Additive, pH 9, adjusted with NH4OH). The gradient program consisted of 0-2 min (96% B), 5.5-8.5 min (88% B), 9-14 min (86% B), 17 min (82% B), 23-24 min (65% B), 24.5-26 min (96% B), and a 10-minute end-run at 96% B. Mass spectrometry was performed using an Agilent 6230 Time of Flight (TOF) mass spectrometer with an Agilent Jet Stream electrospray ionization (ESI) source in negative mode. Data analysis involved peak visualization and confirmation using Profinder 8.0 (Agilent) software and a pathway-specific, manually curated database. Standard metabolites were included in each run for retention time matching and verification. Heatmaps were generated using Prism, with averaged peak heights normalized to the control casamino acids condition.

### Protein Purification

*V. cholerae’s* GlmU gene was amplified from N16961 gDNA and cloned into pET28a downstream of 6xHis-SUMO Tag (61). Plasmids were verified by Sanger sequencing. *E. coli* BL21 (DE3) (Novagen) was transformed with the resulting recombinant plasmid (pET28a-GlmU). Overnight cultures (10 mL) were used to inoculate 1L of LB with kanamycin (50 μg/mL) and incubated at 37°C with vigorous shaking (220 RPM) until they reached an OD600 between 0.6 and 0.8. Cultures were induced with 1mM IPTG at 18°C and 180 RPM overnight. Harvested cells were pelleted and resuspended in 15 mL of cold purification buffer (20mM Tris pH 7.5, 150 mM NaCl), and lysed by sonication. Lysates were cleared by centrifugation at 31,000g for 40 minutes at 4°C, and loaded onto a HisPur cobalt column (Thermo Scientific; Catalog No. 89964) and washed multiple times with purification buffer until protein was undetectable in the flowthrough by the Bradford reagent. The bead slurry was then transferred to a 5 mL microtube with 60 μL of ULP1 Sumo protease and digested overnight at 4°C rotating. Protein was eluted the next day with 20 mL of purification buffer. Samples were analyzed by SDS-page with Coomassie blue stain and then Concentrated with a 30KD Amicon concentrator (Millipore) to 5 mL. Concentrated samples were then measured using Nanodrop.

### *In vitro* GlmU biochemistry

Reaction design was taken from (50). In summary, GlmU reaction substrates included GlcN-1P (5mM), UTP (5mM), and Acetyl-CoA (5mM). substrates were added to a 1.5mL Eppendorf tube with 5μL of 10x Reaction Buffer (50mM Tris–HCl, pH 7.5, 5 and 5mM MgCl_2_). A dilution series of G1P inhibitors were added (0mM – 250mM) and then purified GlmU^VC^ was added at 2 µM, for a total volume of 50μL. The tubes were incubated at 30°C for 30min. Equal volumes of 40:40:20 (Acn: MeOH:H_2_O) solution was added to stop the biochemical reaction. The tubes were then centrifuged for 10min at 15,000rpm. Half of the volume was added to a new tube and mixed with equal volumes of LC-MS Solution B. These tubes were centrifuged at 4°C, 8min, at 15,000rpm. 25μL of supernatant was added to the LC-MS autosampler vials and 2 µl sample volume were resolved on a Diamond Hydride Column using a 1260 Infinity II high-performance liquid chromatography (LC) system (Agilent) coupled with an Agilent Accurate-Mass 6230 TOF-Mass Spectrometer (MS) operating in negative mode. Two liquid phases (i) solvent A: H_2_O + 0.2% formic acid and (ii) solvent B: Acetonitrile + 0.2% formic acid were used at 0.4 ml/ min with the following gradients: 85% B, 0-2min; 80% B, 3-5min; 75% B, 6-7 min; 70% B, 8-9 min; 50% B, 10-11 min; 20% B, 11-14 min; 5% B, 14-24 min and 10 min of 85% B for the re-equilibration. Results were collected on Agilent 6230 TOF-MS with ESI source. Profinder 8.0 (Agilent) was used for the peak abundance measurement. Final metabolites were verified by comparing retention times and mass-to-charge (m/z) ratios with respective standards for each substrate and product. Absolute and relative counts were calculated and plotted on GraphPad Prism software.

### Molecular Modeling

Interactions between GlmU, glucosamine-1-P, and glucose-1-P were modeled using Chai Discovery (https://www.chaidiscovery.com/blog/introducing-chai-1). We input the amino acid sequence code for VCH GlmU, from UniProt (<u>Q9KNH7</u>) as the protein input x3. We then uploaded the SMILES for either <u>glucosamine-1P</u> or <u>glucose-1P</u> (PubChem). Confidence scores (pTM and ipTM) were automatically generated during this analysis. Each resulting model was visualized using PyMol. Polar interactions to adjacent amino acids were identified and measured in PyMol. Pairwise structural alignments were performed in PyMol and RMSD values were automatically generated during this analysis. Sequence alignments and analysis were performed using UniProt (https://www.uniprot.org/align).

## Acknowledgements

MRK is supported by the National Institutes of Health and National Institute of Allergy and Infectious Diseases Award T32AI145821. Tolerance research in the Dörr lab is supported by NIH R01 AI143704. The Rhee lab is funded by NIH R25 AI140472.

## Supporting Information

**Figure S1:**
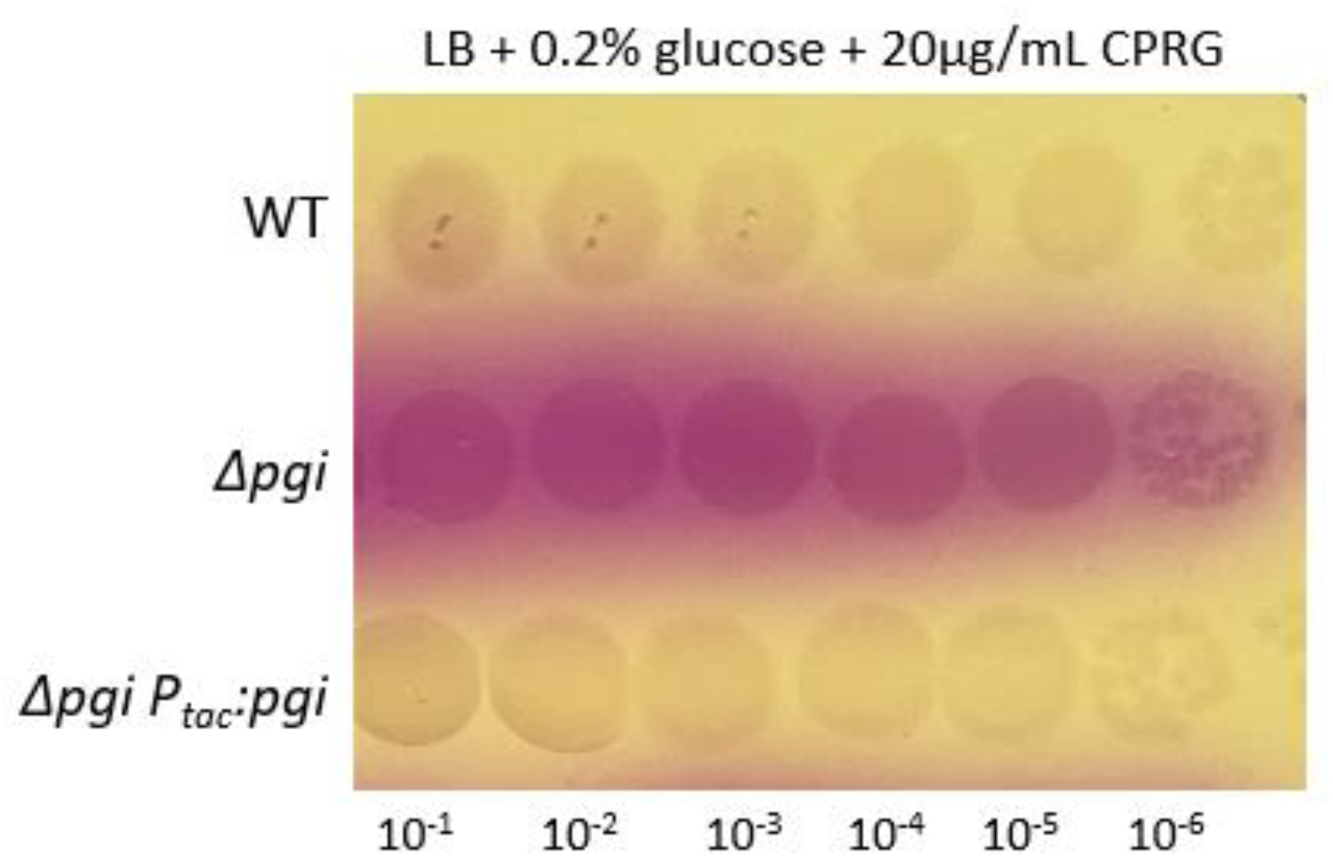
Enhanced lysis of a *pgi* mutant. Overnight cultures of the indicated strains were plated on agar containing glucose and the lysis indicator CPRG (see text for details) and imaged after 18 hours of growth.

**Figure S2:**
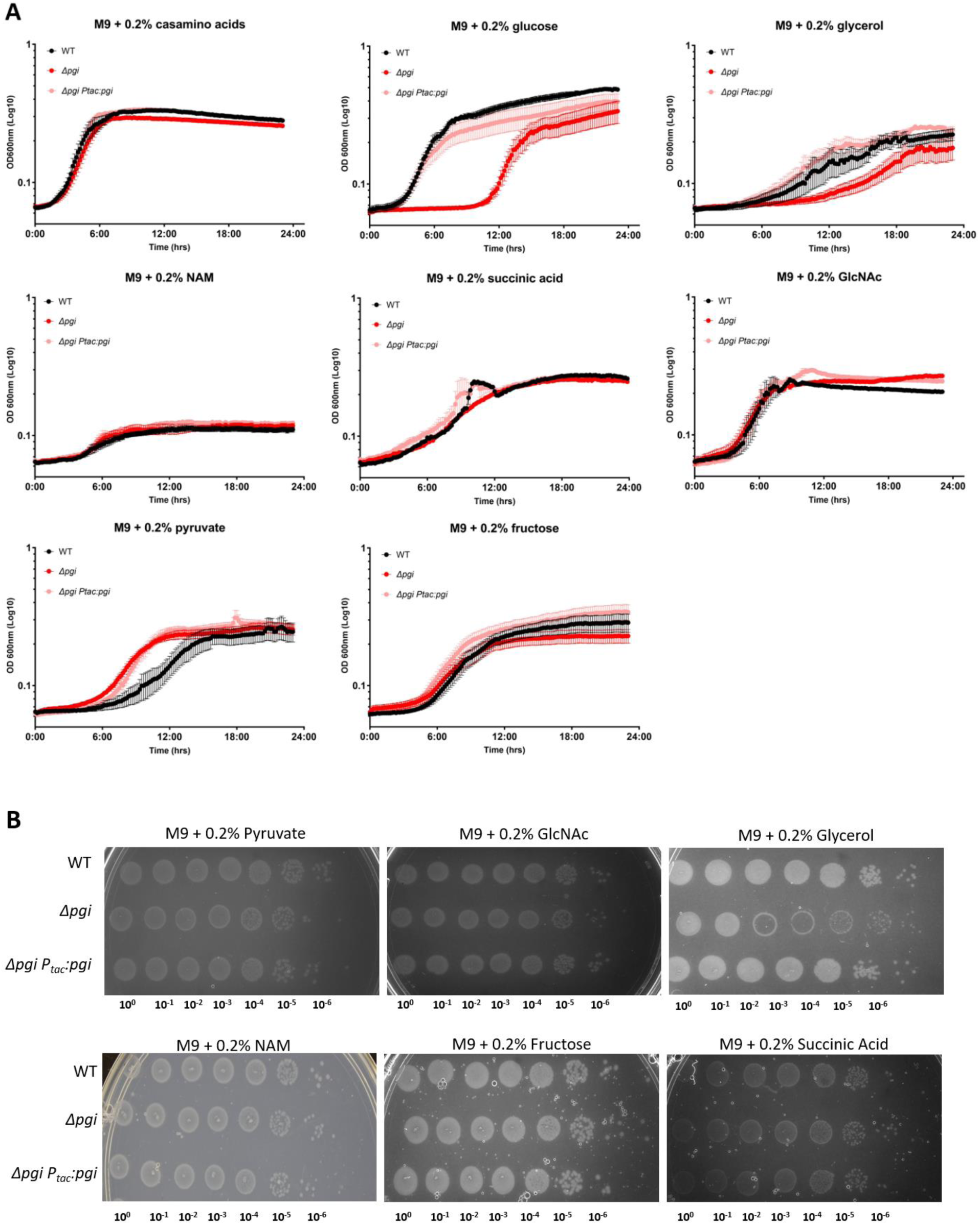
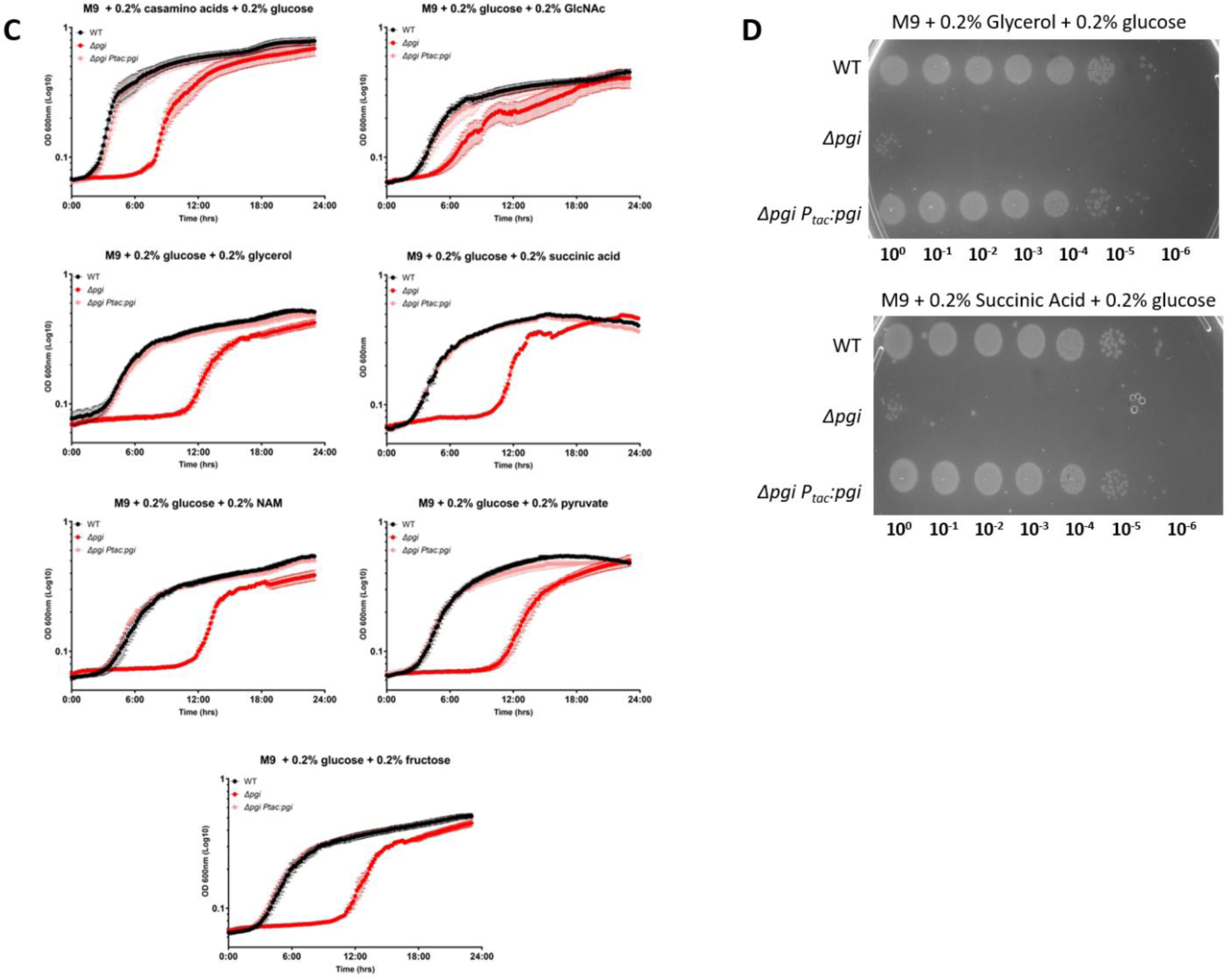
Ability of Δ*pgi* to grow on various carbon sources. A) Growth curves with M9 minimal media and the indicated sole carbon sources. SEM plotted. B) Serial dilutions of overnight cultures grown in M9 + 0.2%CAA plated on M9 agar supplemented with the indicated carbon sources and grown for 18 hours at 37°C. C) Growth curves with M9 minimal media, designated carbon sources, and 0.2% glucose. SEM plotted. D) Serial dilutions of overnight cultures grown in M9 + 0.2%CAA plated on M9 agar supplemented with the designated carbon sources and 0.2% glucose.

**Figure S3:**
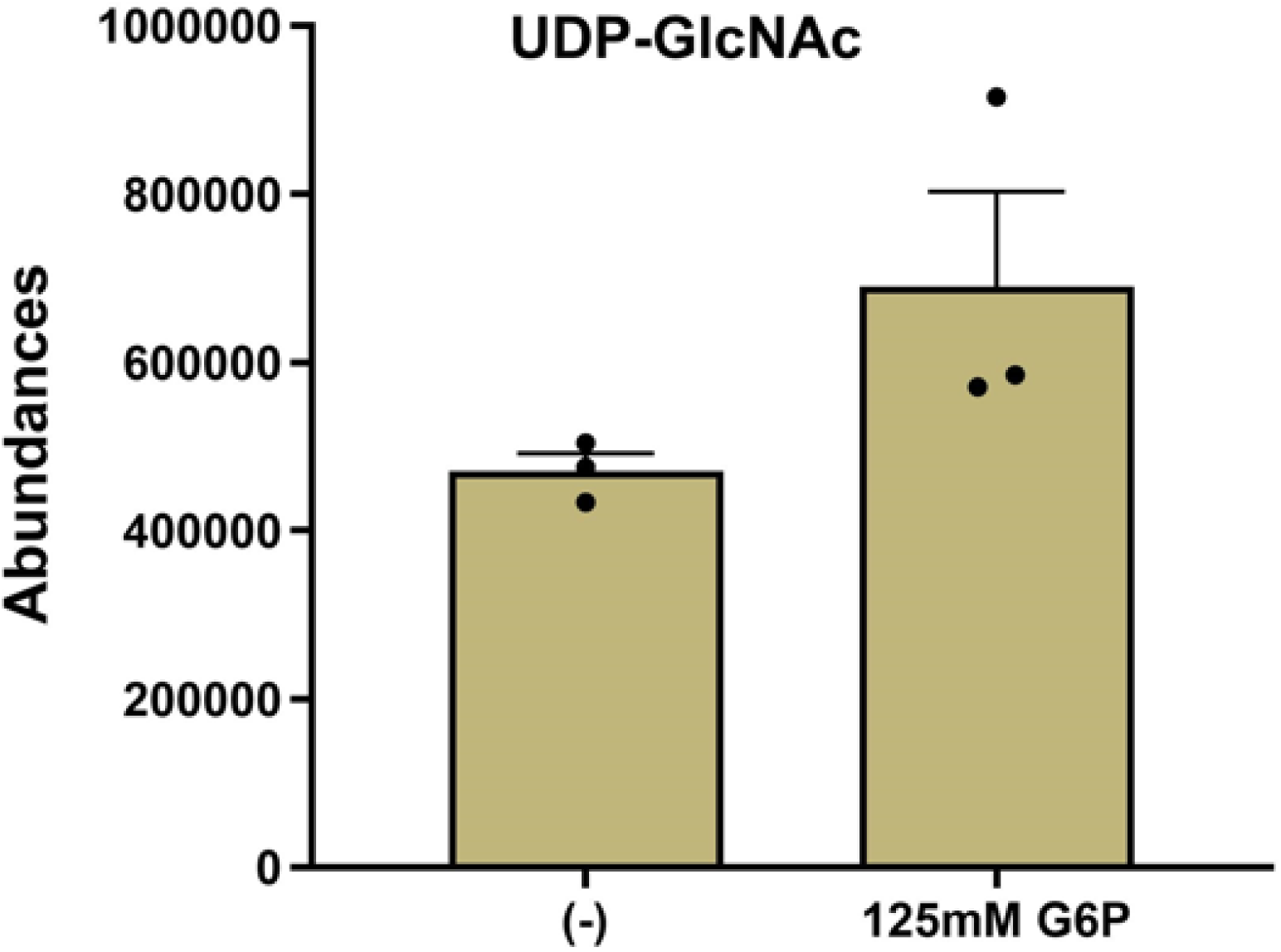
*In vitro* biochemical abundance of specified metabolites. UDP-GlcNAc levels measured with 125mM G1P addition. SEM plotted with 3 replicates displayed.

**Figure S4:**
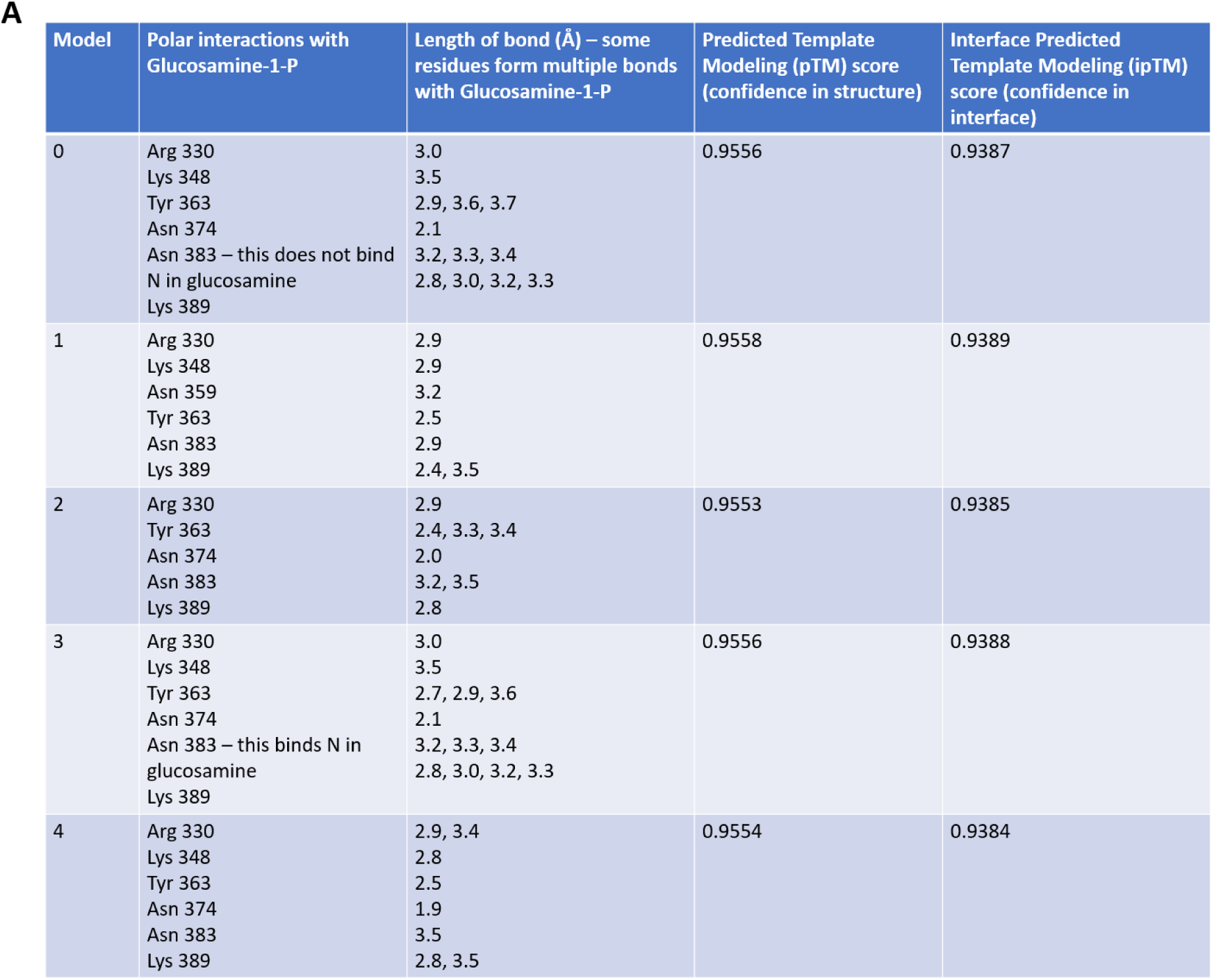
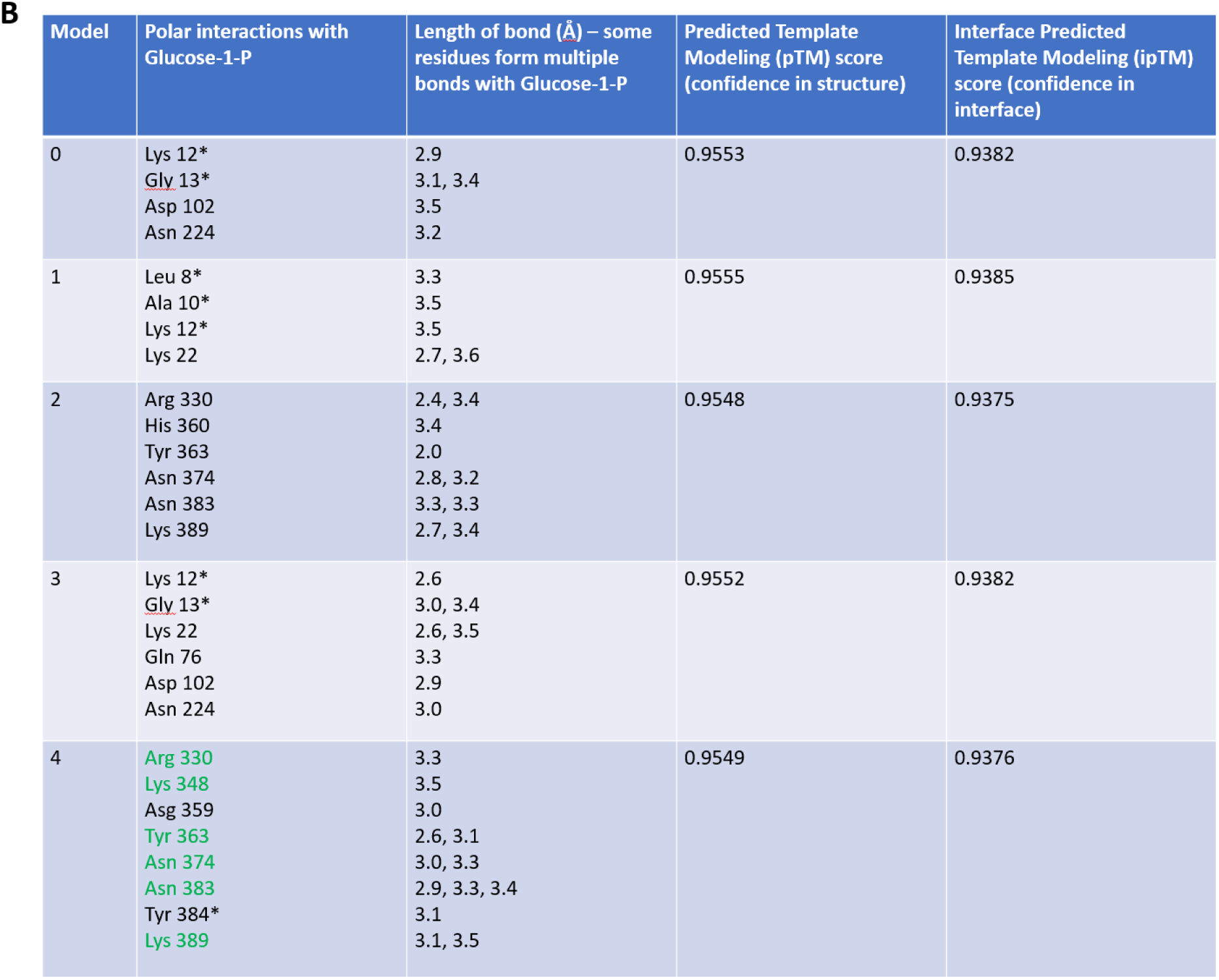
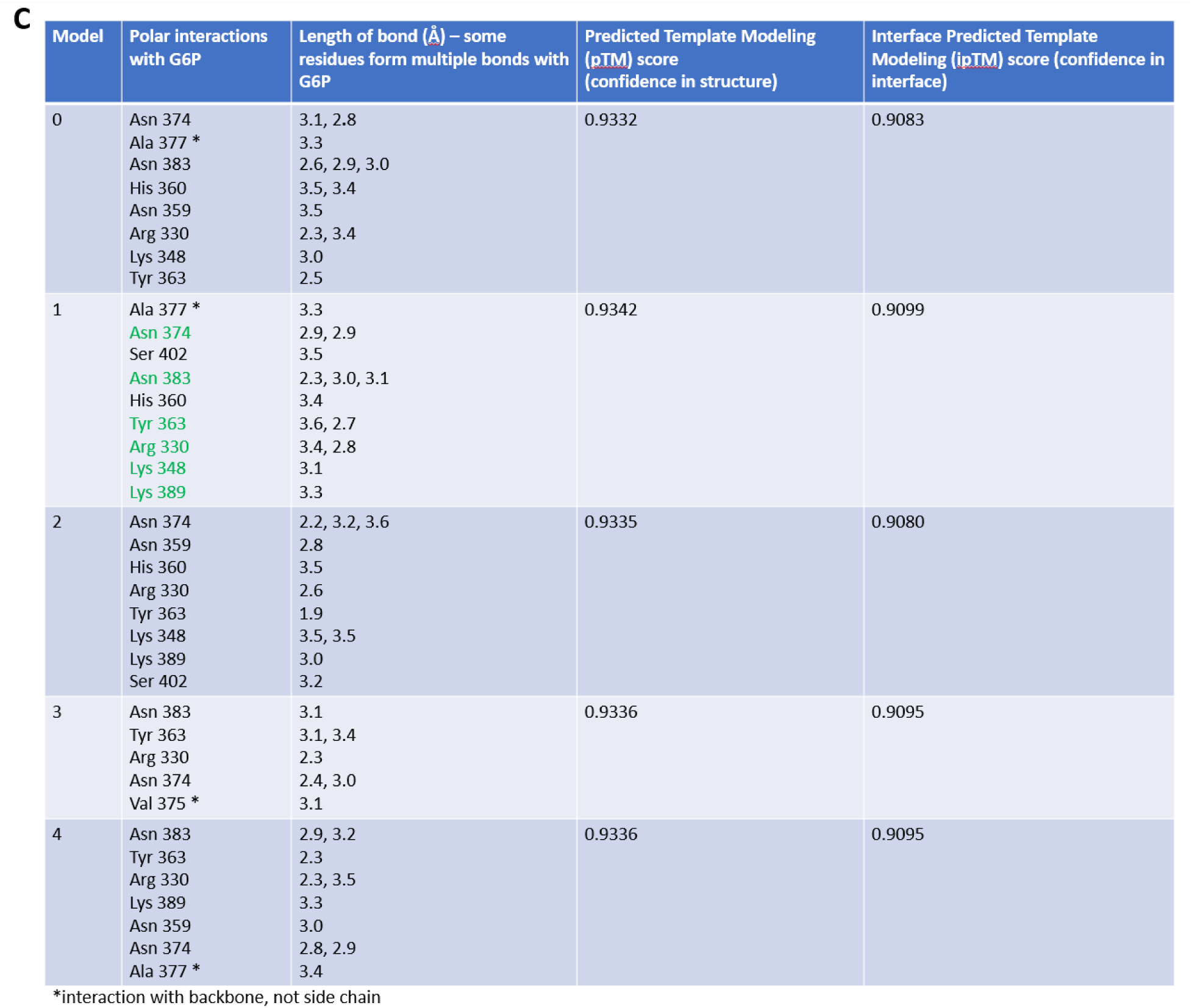
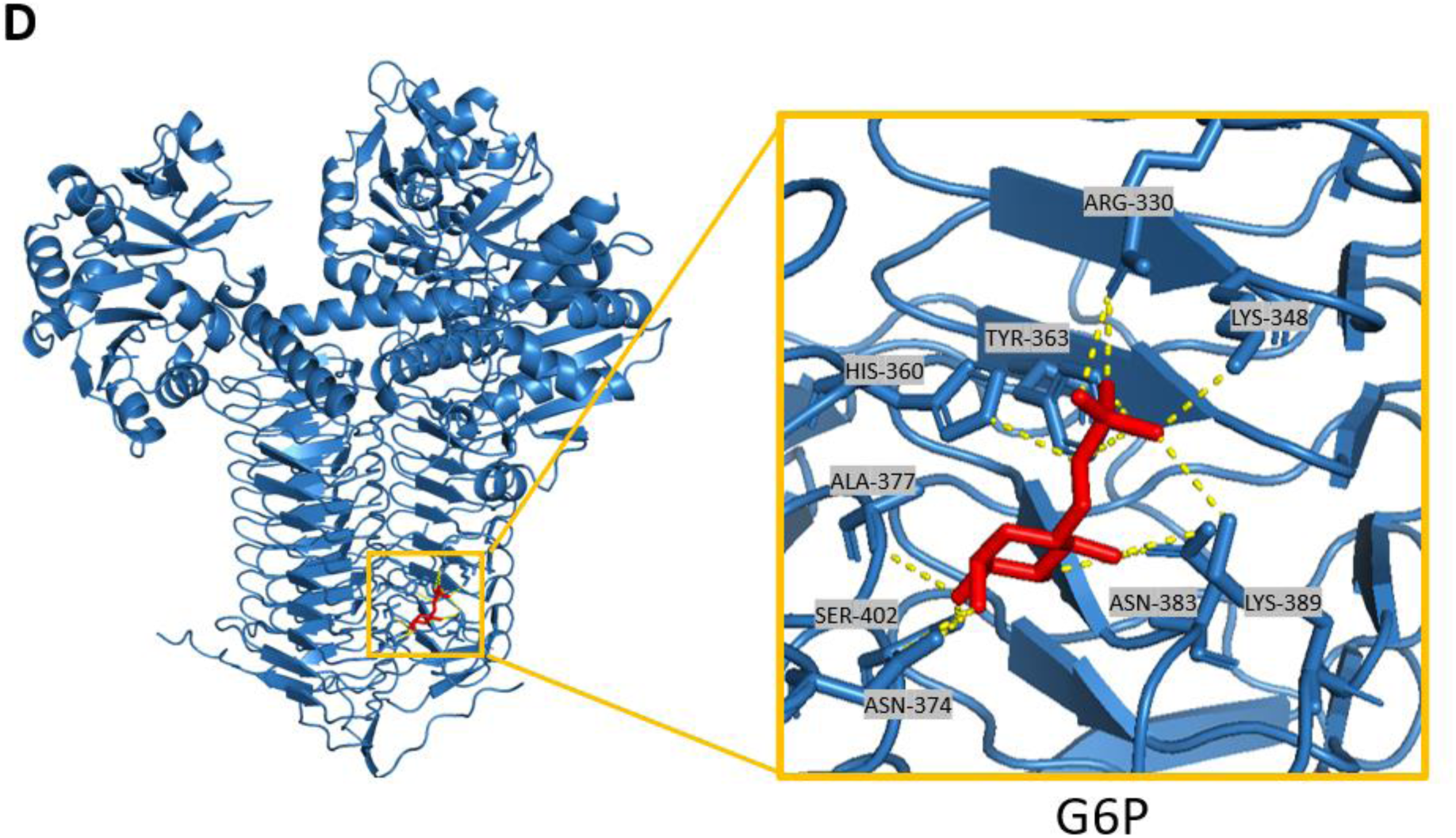
Predicted molecular models of GlmU. A) Predicted models of VCH GlmU with glucosamine-1P with interacting residues denoted, length of bonds, pTM and ipTM scores. Model #3 was the model used above. B) Predicted models of VCH GlmU with glucose-1P with interacting residues denoted, length of bonds, pTM and ipTM scores. Model #4 was the model used above. Highlighted in green are the same interacting residues as the substrate GlcN-1P. C) Predicted models of VCH GlmU with glucose-6P with interacting residues denoted, length of bonds, pTM and ipTM scores. Model #1 was the model used above. Highlighted in green are the same interacting residues as the substrate GlcN-1P. D) Molecular modeling of GlmU binding glucose-6P (red) and the associated polar interactions.

**Figure S5:**
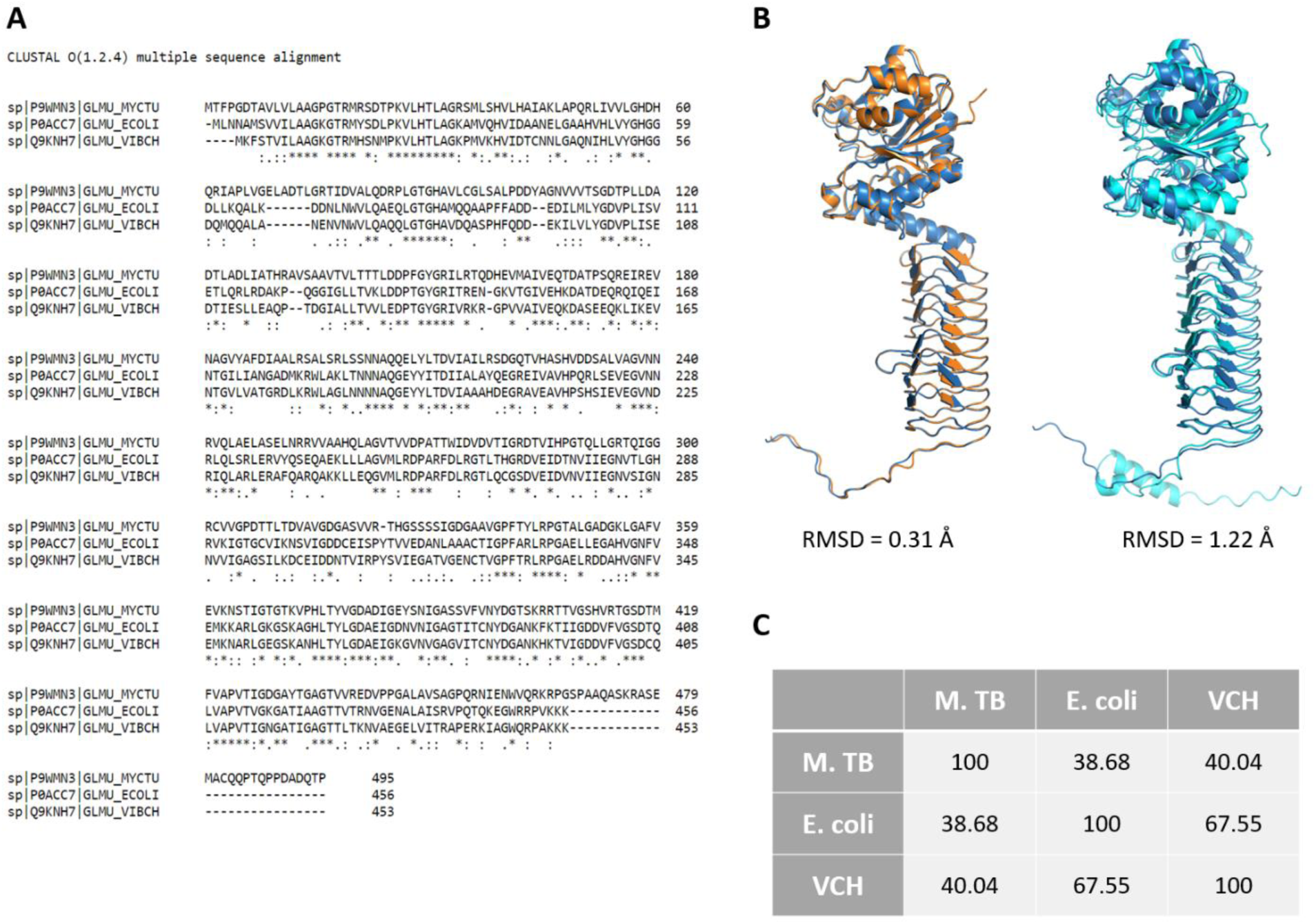
Structural alignments of GlmU. A) Multiple amino-acid sequence alignments of *M.Tb, E.coli,* and *V. cholerae* with conserved residues denoted, using Clustal. B) GlmU monomer structural alignment. GlmU^VC^ is blue, GlmU^EC^ is depicted in orange, and GlmU^Mtb^ is in teal. RMSD, root mean squared deviation, is a measure of how closely two alignments match; RMSD < 2.5 is a reasonable alignment. C) Percent Identity Matrix, created by Clustal2.1 shows alignment similarity across species.

**Table S1:** Strain list used in this study. * = checked via whole genome sequencing.

| Strain | Description | Source or Reference |
| --- | --- | --- |
| WT* | Wild-type *V. cholerae* N16961 El Tor | Heidelberg JF, *et. al.*, 2000 |
| MK1* | $\Delta pgi$ (vc0374) | Keller MK, *et. al.*, 2023 |
| MK4 | $\Delta pgi$ P<sub>tac</sub>- *pgi* | Keller MK, *et. al.*, 2023 |
| MK12 | $\Delta pgi$ P<sub>tac</sub>- *nagB* (vca1025) | This study |
| MK13* | $\Delta pgi$ $\Delta pgcA$ (vc2095) | This study |
| MK14 | $\Delta pgi$ P<sub>tac</sub>- *pgcA* | This study |
| MK15 | $\Delta pgi$ P<sub>tac</sub>- *glmS* (vc0487) | This study |
| MK16 | $\Delta pgi$ pHL: *glmU* (vc2762) | This study |
| MK110 | *E. coli* SM10 λpir conjugation strain | Ferrières L, *et. al.*, 2010 |
| MK120 | *E. coli* MFD pir conjugation strain | Ferrières L, *et. al.*, 2010 |
| MK130 | *E. coli* BL21 protein purification strain | Novagen |

\* = checked via whole genome sequencing

**Table S2:**
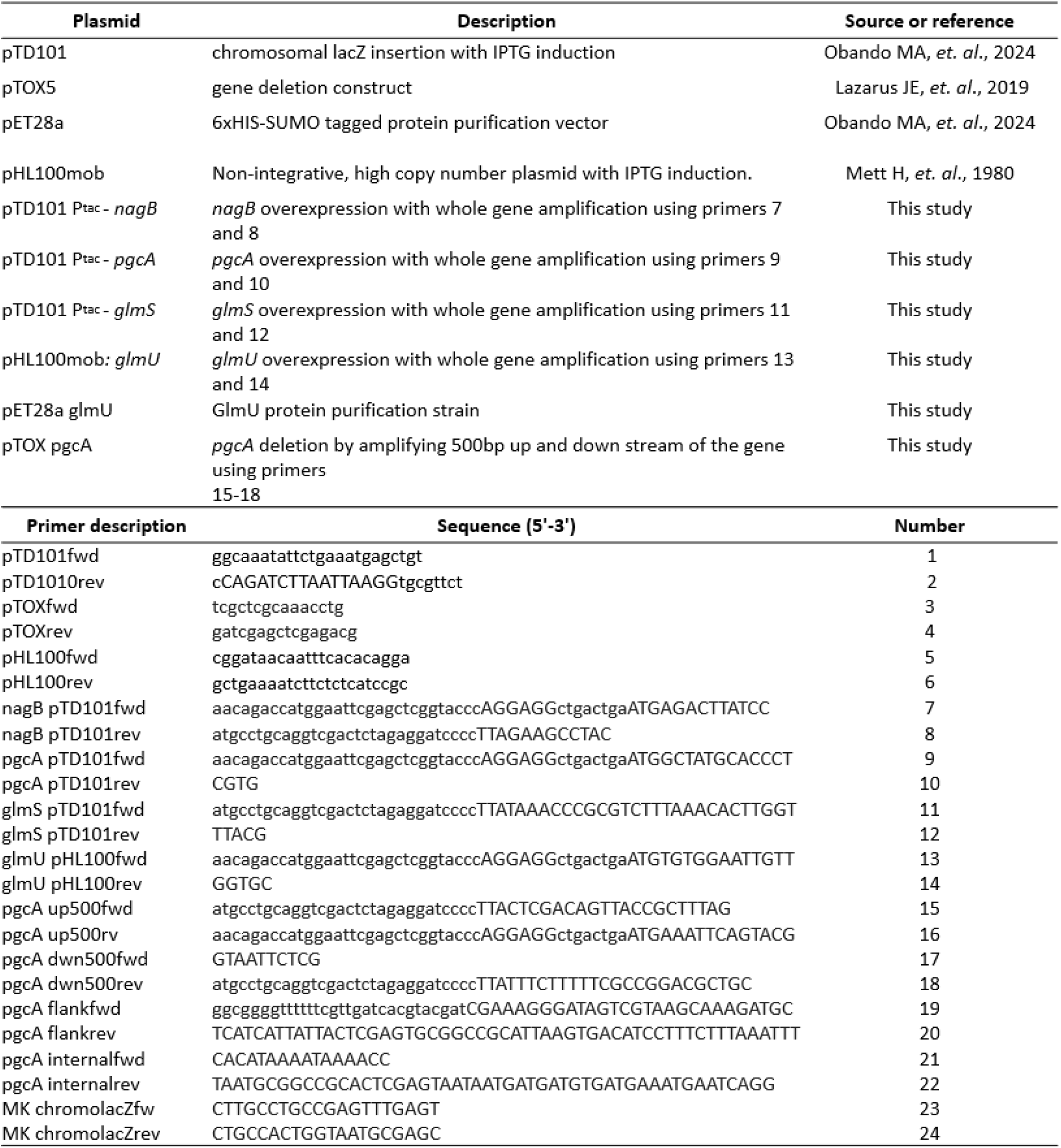
Oligos used in this study.

